# TMCC2 Associates with Alzheimer’s Disease Pathology in the Human Brain

**DOI:** 10.1101/2021.08.14.456332

**Authors:** Paul C. R. Hopkins, Claire Troakes, Andrew King, Guy Tear

## Abstract

Transmembrane and Coiled-Coil 2 (TMCC2) is a protein that forms complexes with both apolipoprotein E (apoE) and the amyloid protein precursor (APP), two proteins central to the etiology of Alzheimer’s disease (AD). Here we have carried out the first investigation of TMCC2 expression in the human brain, where TMCC2 immunoreactivity is primarily neuronal. We also examined TMCC2 localization in late onset AD brains,early onset AD cases bearing a mutation in APP, and in Down syndrome cases. TMCC2 immunoreactivity is closely associated with APP immunoreactivity in both control and AD cases. In all AD cases examined, TMCC2 associated with dense core senile plaques and adjacent dystrophic neurites, but not with amyloid surrounding the core, nor with diffuse amyloid plaques or neurofibrillary tangles. In Down syndrome, we found dystrophic neurites in senile plaques that were positive for amyloid and TMCC2, but not for APP, in addition to the APP- and TMCC2-positive dense cores. Western blot analysis revealed TMCC2 exists as at least three protein isoforms. The relative abundance of the isoforms varies in the temporal gyrus and cerebellum, the relative proportions of which were influenced by APOE and/or dementia status.

Taken together, these observations are consistent with the evolutionarily conserved interactions between APP and TMCC2 and suggest cooperation betweenTMCC2 and APP in the normal and AD brain. These findings thus support a role for TMCC2 as a mediator of the impact of apoE on AD pathogenesis.

## Introduction

The ε4 isoform of apolipoprotein E (apoE4) is the greatest risk factor for late onset AD after age (Genin et al, 2011; Raber et al, 2004). ApoE4 alters the metabolism of Amyloid Protein Precursor (APP), and/or its derivative, the Aβ peptide, more than the more common apoE3 isoform, leading to an increase in amyloid plaques in both human tissue analysis studies and experimental mice, (Holtzman et al, 1999; Ossenkopple et al, 2015). Yet the mechanisms linking apoE isoforms, APP and/or Aβ metabolism, and risk of neurodegeneration are poorly understood.

Transmembrane and Coiled Coil 2 (TMCC2) is a protein that forms complexes with APP and apoE (Hopkins et al, 2011) suggesting it could mediate an influence of apoE on APP processing. In the presence of apoE, TMCC2 mediates an increase in Aβ secretion from cells expressing an autosomal dominant mutated form of APP, APPswedish (K595N, M596L), and from the C-terminal fragment of APP that is generated in vivo following cleavage by BACE. The AD associated apoE4 isoform shows a larger effect on Aβ production than apoE3 in the presence of TMCC2 (Hopkins et al., 2011). TMCC2 thus shows differential interactions with normal versus risk forms of two separate proteins implicated in the genetic etiology of AD.

The interaction of TMCC2 with APP is evolutionarily conserved: its *Drosophila* orthologue, called Dementin, is highly similar to TMCC2 in amino acid sequence and protects against ectopic expression of human APP *in vivo* (Hopkins, 2013). When mutated, Dementin perturbs the metabolism of the endogenous *Drosophila* orthologue of APP, the APP-Like protein, and causes neurodegeneration with pathological features of AD, including accumulation of fragments of the APP-Like protein, synaptic defects and early death (Hopkins, 2013).

Here we report the first investigation of TMCC2 expression in the human brain. We also examined TMCC2 distribution in a variety of AD patients; late onset AD patients stratified by apoE genotype and age-matched non-demented controls, early onset AD associated with mutations in APP or Down’s syndrome AD. We found that TMCC2 is affected by both APOE genotype and brain region and associates with hallmark pathologies of AD in both early and late onset AD, implicating this protein in AD pathogenesis.

## Results

### TMCC2 in human brain

Using a rabbit antibody previously validated against mouse and rat brain-derived TMCC2, (antibody 94) (Hopkins et al, 2011), we detected a set of three bands in human brain homogenates by western blot. These migrated at a position close to that predicted from the amino acid composition of TMCC2 (77.5 kDa; Fig 1A). A similar pattern was observed using a separate independently generated antibody (antibody 11193), but which had reduced specificity compared to antibody 94 (Fig. S1A).

**Figure 1.**
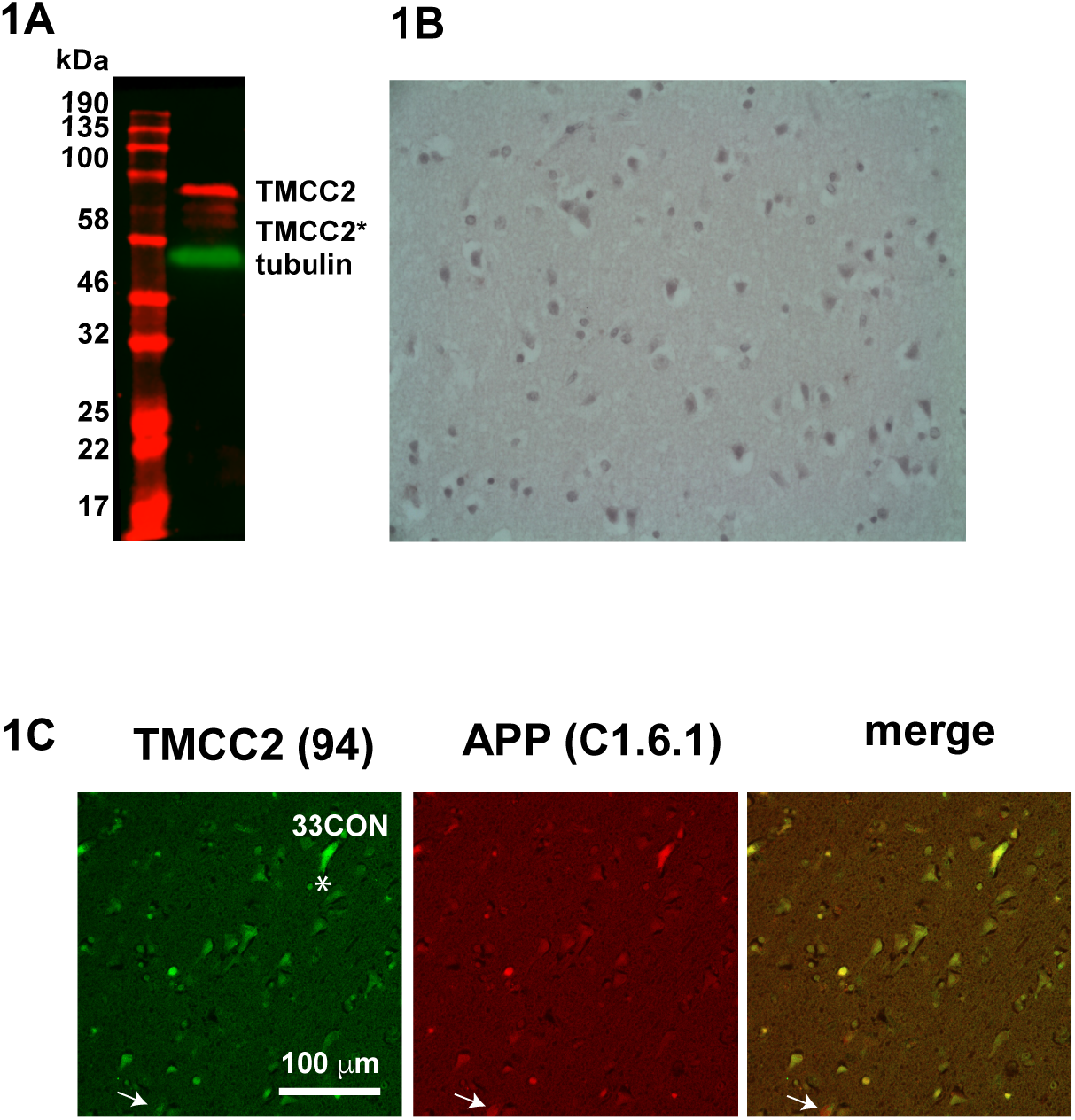
Expression of TMCC2 in the human brain detected using antibody 94. **1A**, western blot of human superior temporal gyrus for TMCC2; **1B**, immunohistochemical anayysis for TMCC2 in human superior temporal gyrus of an APOE3 homozygous non-demented case; **1C**, dual immunofluorescent detection of TMCC2 (green) and APP (antibody C1/6.1, red) in a non-demented APOE3 homozygous temporal gyrus, arrow in 1C indicates rare staining for APP that is not co-incident with staining for TMCC2, the asterisk indicates a blood vessel staining for both TMCC2 and APP.

Immunohistochemical staining for TMCC2 in the temporal cortex showed primarily a neuronal somatodendritic pattern (Fig. 1B), consistent with previous detection in primary neurons (Hopkins et al., 2011) and proposed roles of TMCC2 in endomembrane systems (Hopkins et al, 2011; Hoyer et al., 2018). TMCC2 forms a complex with APP in transfected human cells, and in rat brain, where a large fraction of TMCC2 exists in a complex with APP (Hopkins et al, 2011). We therefore examined human brain sections for both TMCC2 and APP by double immunofluorescence using antibody 94 to TMCC2 and antibody C1/6.1, which binds the C-terminus of APP (residues 676–695 of APP_695_) (Mathews et al, 2002). We found a high degree of similarity between the staining patterns for both (Fig. 1C). In cognitively healthy brains, TMCC2 immunoreactivity was rarely distinct from that for APP, while APP staining exists either together with or separate from TMCC2 (Fig 1C, arrow), suggesting a dynamic association between TMCC2 and APP. TMCC2 and APP immunoreactivity was also detected in the lumen of blood vessels (Fig 1C, asterisk).

### TMCC2 in late onset Alzheimer’s disease

To investigate TMCC2 in late onset AD, we examined TMCC2 protein levels and cellular distribution in five APOE3 homozygous cognitively healthy human brains with an average age of 85 (range 82-90), five AD cases homozygous for APOE3 pathologically diagnosed with AD (Braak stages IV-V) with an average age of 77 (range 70-81), and five AD cases homozygous for APOE4 and diagnosed with AD Braak stages V-VI, average age 76 (range 71-89) (case details are provided in supplementary Table 1).

In late onset AD brains from both APOE3 and APOE4 homozygotes, neurons had a shrunken appearance, with TMCC2 immunoreactivity distributed similarly to that seen in age-matched non-demented controls (Fig 2A and B). To investigate whether TMCC2 associates with amyloid plaques we stained AD cases using anti-TMCC2 antibody 94 and anti-Aβ (antibody 4G8). TMCC2 immunostaining was notable in the centre of dense core senile plaques from cases homozygous for either APOE allele but did not specifically associate with the diffuse amyloid halo surrounding dense cores (Figs. 2C and 2D, and further figures below), or in diffuse plaques without a dense core (Fig. 2E and further figures below). Specificity of TMCC2 detection in these plaques was confirmed using the separate antibody 11193 (Fig. S2A).

**Figure 2.**
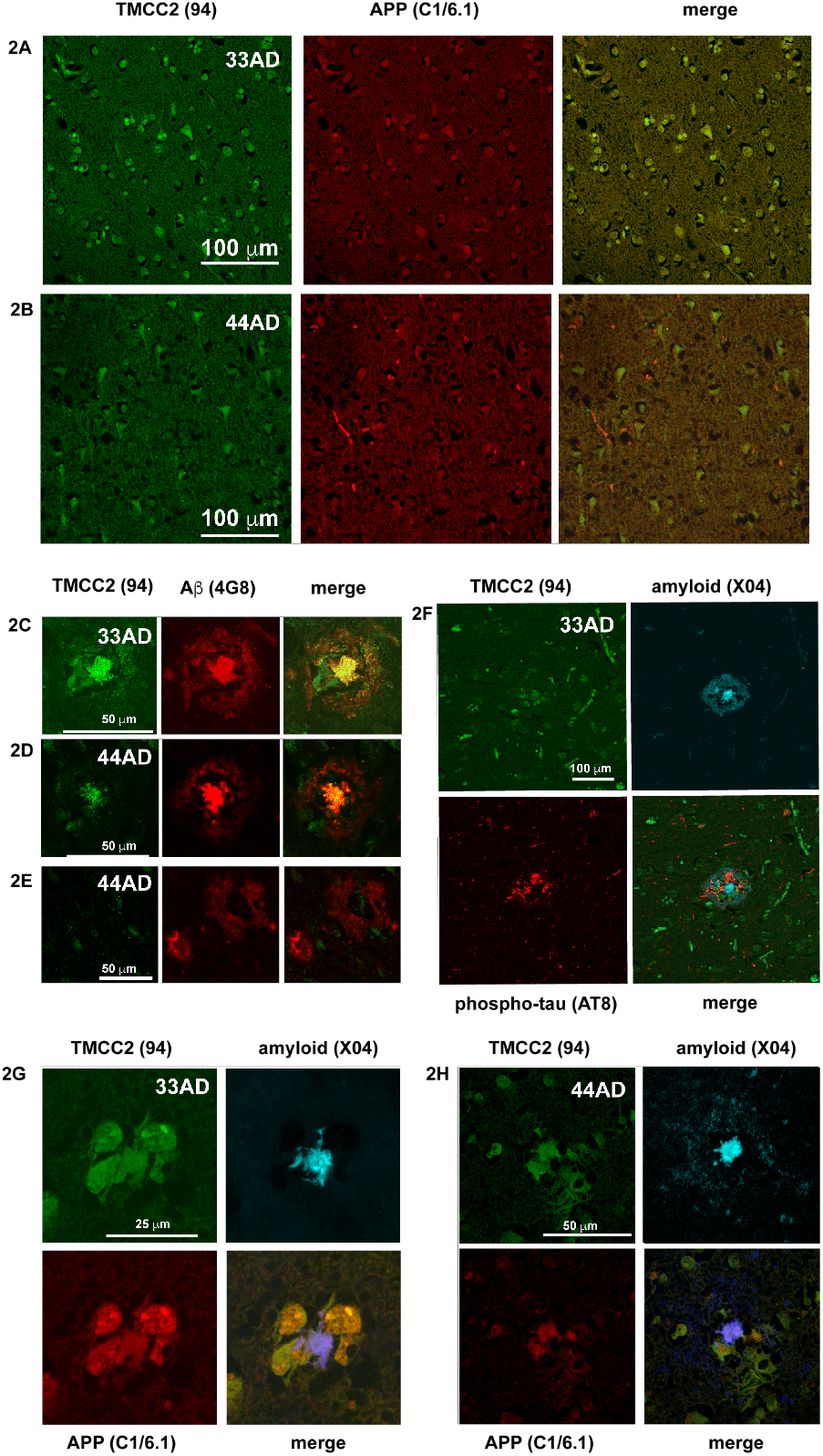
Association of TMCC2 with late onset AD pathology in human superior temporal gyrus. TMCC2 immunoreactivity was co-incident with APP immunoreactivity in the temporal gyrus of both APOE3 homozygotes (**2A**) and APOE4 homozygotes (**2B**) with AD. TMCC2 was detected in dense cored amyloid plaques of APOE3 homozygotes and APOE4 homozygotes detected with antibody 4G8 (**2C** and **2D**), but not in diffuse amyloid plaques (**2E**). TMCC2 was not detected in phospho-tau-positive dystrophic neurites (**2F**). TMCC2 and APP immunoreactivity co-localized in APP-positive dystrophic neurites surrounding dense cored plaques detected with methoxy-X04 in both APOE3 homozygotes and APOE4 homozygotes with AD (**2G** and **2H**).

We next examined if TMCC2 might associate with phospho-tau-filled neurites detected by antibody AT8, (which recognises tau phosphorylated at both serine 202 and threonine 205) surrounding dense cored amyloid plaques detected using methoxy-X04 (a derivative of Congo Red that labels amyloid). We found no notable association of TMCC2 and AT8 immunoreactivity in AT8-positive neurites (Fig 2F).

Amyloid plaques of AD present in multiple morphologically and immunochemically distinct types. Among plaques having dense amyloid-positive cores a subset may contain phospholipid shells (Rak et al, 2007) and also stain for cellular proteins (D’Andrea & Nagele, 2010; Pensalfini et al, 2014), thus may derive in part from degenerating neurons. We examined the distribution of TMCC2, APP and amyloid in late onset AD brain sections homozygous for either APOE3 or APOE4. This detected presumptive dystrophic neurites immunopositive for both APP and TMCC2 adjacent to dense amyloid cores detected using methoxy-X04 (Figs. 2G and 2H). The vast majority of such dense-core-associated APP-positive dystrophic neurites co-stained for both APP and TMCC2; separate staining occurred in only 2.5% of such plaques imaged across both APOE genotypes (e.g. supplementary Fig. S2B).

### TMCC2 in early onset Alzheimer’s disease

The above findings suggested an association of TMCC2 with late onset AD pathology. We extended our investigation to examine TMCC2 distribution in other examples of AD and examined early onset AD caused by mutation or over-expression of APP, five cases of early onset AD associated with the APP Val717 to Ile mutation (APP London), and one case caused by mutation of APP Val717 to Gly, as well as in three cases of dementia with Down Syndrome where the gene for APP is present in excess.

In early onset AD cases with APP-Val717Ile, immunostaining of temporal gyrus for TMCC2, APP and amyloid, showed extensive amyloid deposits and dense core plaques (Fig 3A). Dense core plaques where the surrounding dystrophic neurites stained for both APP and TMCC2 (Fig 3A’) were found, as were senile plaques surrounded by dystrophic neurites that stained for APP but not TMCC2 (Fig. 3A’’). However, distinct from late onset AD, we also found dystrophic neurites that were positive for APP but not TMCC2 (arrowheads in 3B and 3B’). Similar observations were made for the APP-Val717Gly case (Figs. 3B, 3B’ and 3B’’).

**Figure 3.**
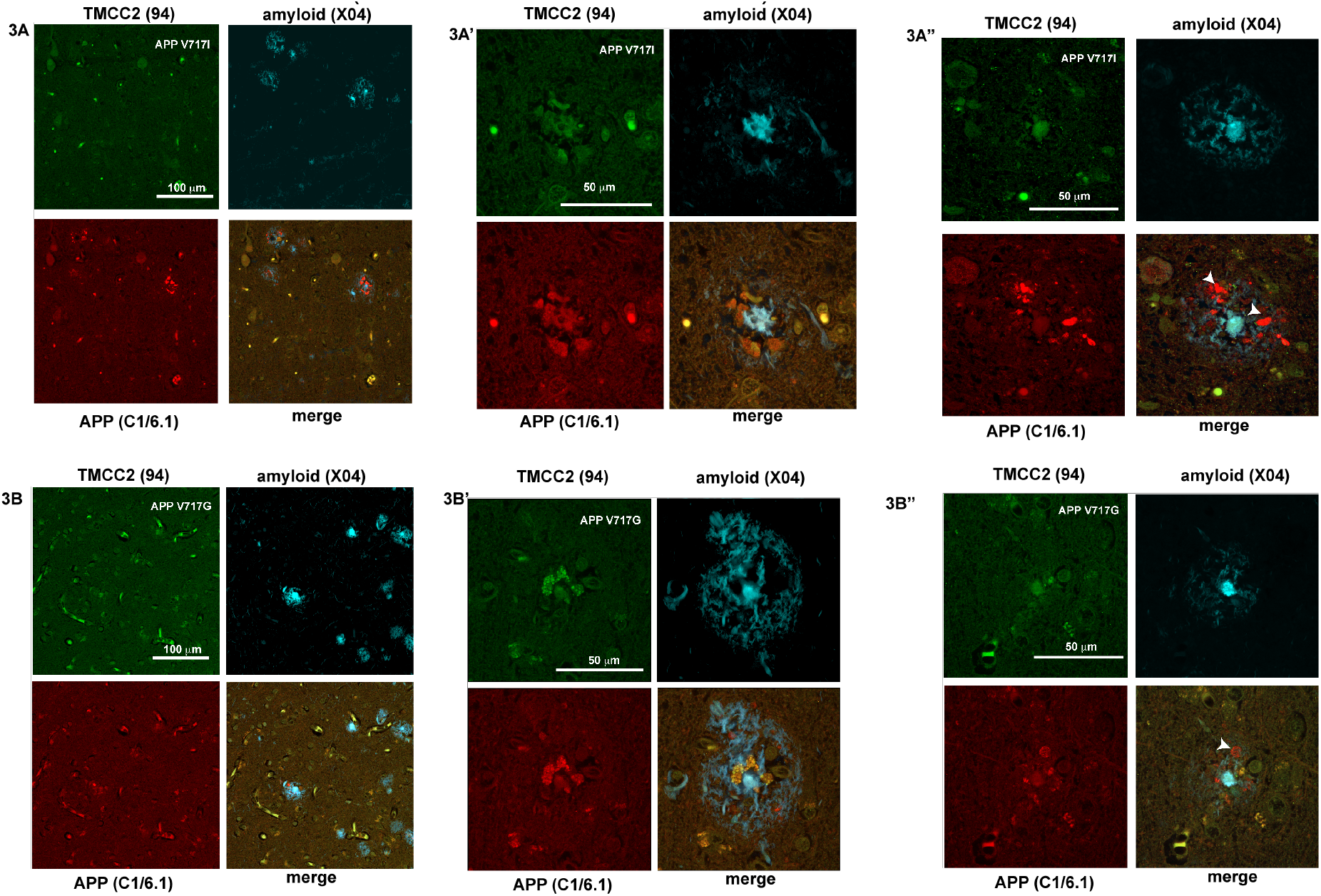
TMCC2 in familial early onset AD. TMCC2 associates with dense cored amyloid plaques and adjacent APP-positive dystrophic neurites in temporal gyrus of familial AD. **3A** and **3B**, abundant amyloid plaques in the temporal gyrus of an APP V717I case and an APP V717G case, respectively, detected with methoxy-X04. Dystrophic neurites surrounding dense cored plaques resembling those found in late onset AD and which stained for both TMCC2 and APP were found (**3A’** and **3B’**), as well as APP-positive dystrophic neurites that did not stain for TMCC2 (arrowheads in **3A’’** and **3B’’**).

We investigated TMCC2 in three Down syndrome cases with a post-mortem pathological diagnosis of AD (details in supplementary Table 1). In all cases we detected abundant amyloid using methoxy-X04, and dense cored plaques that were associated with dystrophic neurites. Both these sites of AD pathology contained both APP and TMCC2 with a distribution similar to those found in familial and late onset AD (Fig. 4A and 4A’). Similar to familial AD and unlike late onset AD, we noted dystrophic neurites enriched for APP but not TMCC2 (arrowheads in 4A and 4B’). In two of the three Down syndrome cases examined, we additionally identified TMCC2 in putative neuritic structures having a spicular or threadlike appearance within senile plaques. These features also contained amyloid as detected by methoxy-X04, but did not contain APP (Figs. 4B and 4B’).

**Figure 4.**
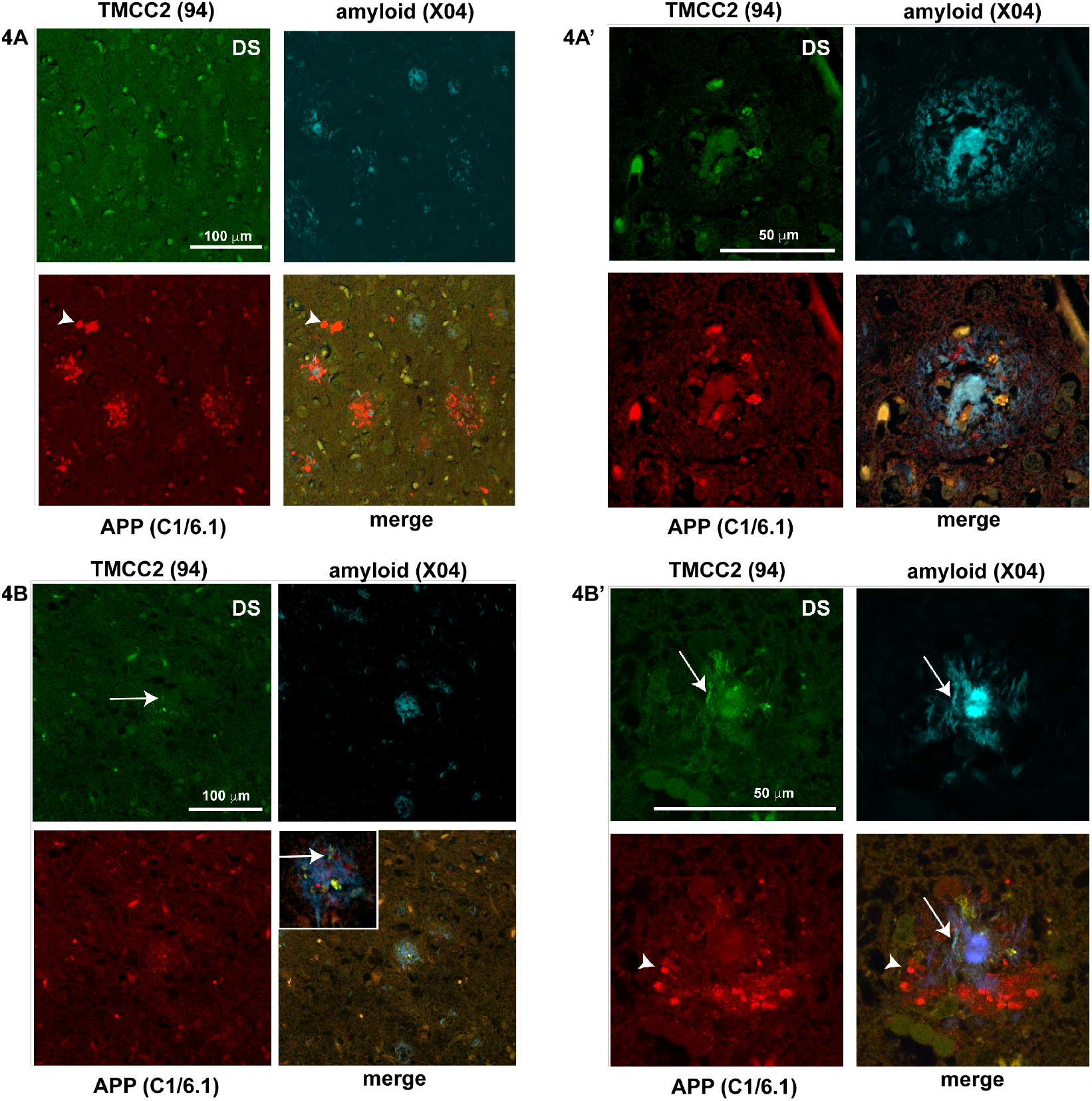
TMCC2 in Down syndrome AD. **4A**, Abundant amyloid plaques and APP-positive dystrophic neurites in Down syndrome temporal gyrus with a pathological diagnosis of AD. **4A’** amyloid plaques resembling those found in late onset AD, i.e. with a dense core detected with methoxy-X04 with adjacent dystrophic neurites staining for both APP and TMCC2 were found in all 3 cases. In all cases, frequent dystrophic neurites that stained for APP but not TMCC2 were seen (arrowheads in **4A** and **4B’**). In two of three cases, dystrophic neurites that were enriched for TMCC2 and amyloid but not APP were identified (arrows in **4B** and **4B’**).

### TMCC2 isoform distribution varies by brain region

Western blot for TMCC2 shows three bands that varied in relative intensity (Figs 1 and 5), which potentially correspond to alternate forms of TMCC2. We examined whether the prevalence of the different TMCC2 isoforms varies or remains constant by brain region, AD or apoE genotype. The total amount of TMCC2 within samples from temporal gyrus or cerebellum were similar. The total levels of TMCC2 was also unaffected by APOE or AD status in the cases assessed (Fig. S5A and S5B). Across all 30 late onset AD and control samples, densitometric quantification of band intensities showed that 57.5 ± 2.7% of TMCC2 in the cerebellum was present in the lower migrating form(s), compared to 42.5 ± 3% in the temporal gyrus (p=0.002 by two-tailed Student’s T-test). This differential expression was statistically significant in APOE3 homozygotes but not in APOE4 homozygotes (Fig. 5C, full blots are shown in Figs. S5A and S5B).

**Figure 5.**
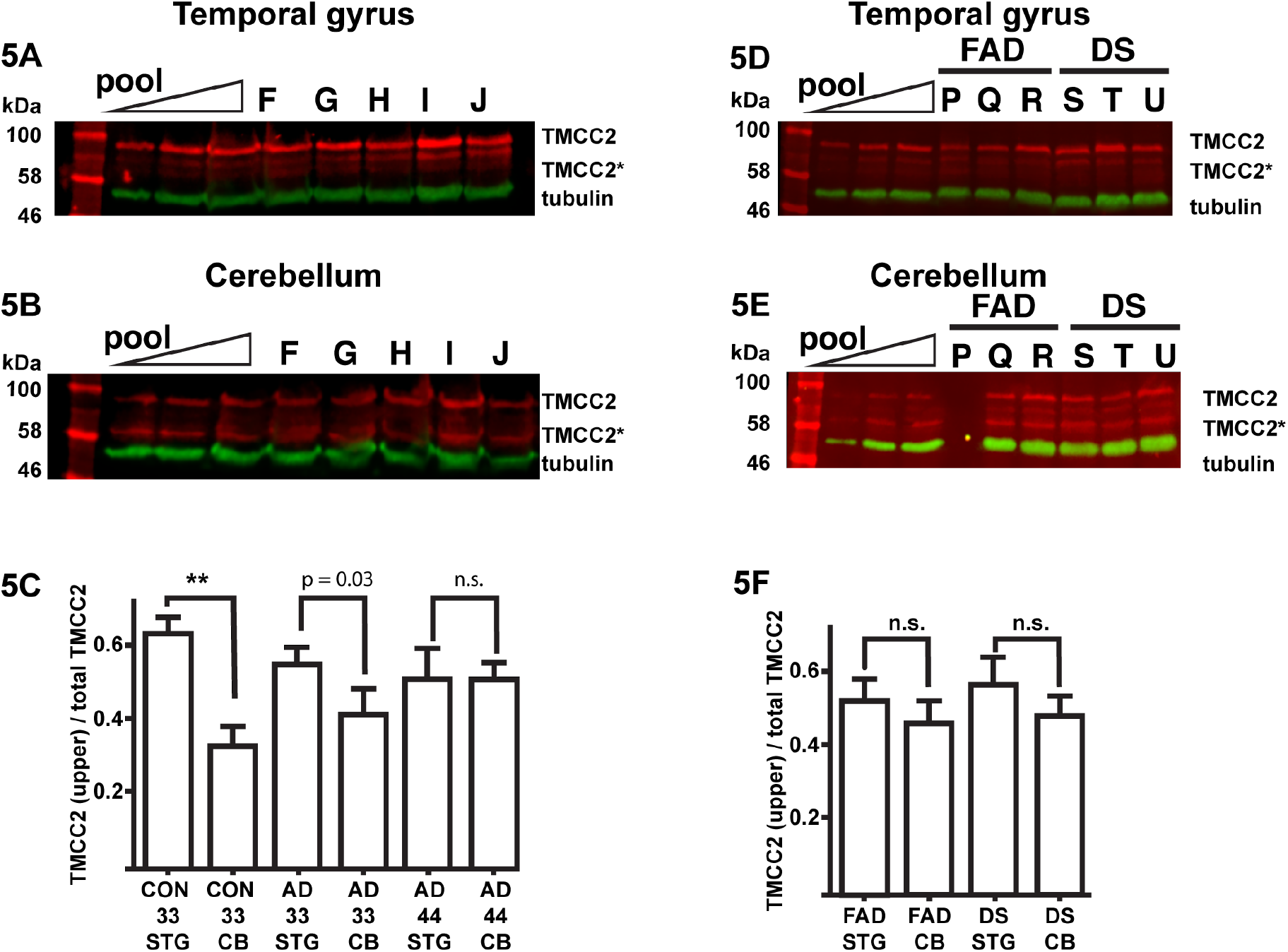
TMCC2 isoforms vary by brain region. Representative western blot of human superior temporal gyrus (**5A)** or cerebellum (**5B**) for TMCC2 in late onset AD and matched controls detected using antibody 94; full blots are shown in Supplementary Figure 5. **5C**, quantification of the ratio of TMCC2 (upper band) to total TMCC2 in brain specimens stratified by brain region, APOE genotype and AD status, statistical significance of differences was estimated by 1 way ANOVA and Tukey’s multiple comparison test (**, p<0.005), or Student’s t-test (p=0.03). **5D** and **5E**, western blot for TMCC2 in temporal gyrus and cerebellum in early onset AD associated with mutation of APP717 (FAD) or Down syndrome (DS). **5F**, no significant differences in the ratios of TMCC2 (upper band) to total TMCC2 between cerebellum (CB) and temporal gyrus (STG) was observed in these samples of early onset AD. Letters over lanes refer to cases described in supplementary Table 1.

Frozen tissue was available from a subset of early onset AD cases and these were analysed as above. This showed a similar variation in the migration of TMCC2 between the temporal lobe and cerebellum as in late onset AD cases and controls (Fig. 5D and E, full blots are shown in Fig. S5C and D). However, the differences did not reach statistical significance (Fig 5F), potentially due to sample numbers being underpowered.

Levels of putative post-translational modification of TMCC2 are therefore potentially influenced by a higher rate of conversion between isoforms in APOE3 homozygous brains, or a reduced conversion in dementia. Thus, the forms of TMCC2 present in the human brain differ according to brain region, as well as APOE status.

## Discussion

TMCC2 has been identified as a molecule which may link apoE status to APP processing and dementia since it complexes with APP, has a differential affinity for apoE isoforms and its Drosophila orthologue can drive neurodegeneration in vivo (Hopkins, et al, 2011; Hopkins, 2013).

In this first study of TMCC2 in the human brain we observed TMCC2 expression within neurons. TMCC2 immunoreactivity showed an association in vivo with that of APP, complementing previous observations of TMCC2-APP complexes in the rat brain and human cells (Hopkins, et al, 2011). An in vivo association of TMCC protein family members with APP protein family members has also been observed in *Drosophila*. The fly orthologue of TMCC2, Dementin, protects against ectopic expression of human APP, and deficiency or mutation of Dementin disrupts processing of the Drosophila APP-Like protein (Hopkins, 2013). The evolutionarily conserved nature of this interaction indicates that the physiological functions of TMCC2 and APP are expected to overlap. TMCC family members are understudied compared to APP family members, nevertheless, emerging data point to converging roles in brain development, neuronal synapse homeostasis (Hopkins, 2013; Heber et al, 2000), and within endomembrane systems (Hopkins et al, 2011; Hoyer et al, 2018).

Given the association of TMCC2 with APP and apoE, and the established role of APP in AD pathogenesis, we investigated TMCC2 distribution in 19 AD cases with varying origins to identify if TMCC2 is associated with AD pathology. We examined late onset AD individuals homozygous for APOE3 or APOE4 together with five age-matched controls, together with early onset AD associated with Down syndrome (three cases) or familial AD associated with APP mutated at Val717 (six cases).

We found a large degree of concordance between the staining patterns for APP and TMCC2 in late onset AD brains and matched healthy controls, where TMCC2 and APP were co-localized with neurons. In all cases of both early and late onset AD brains dense-cored plaques and plaques with dystrophic neurites positive for both TMCC2 and APP were found.

Qualitative differences in staining between early and late onset AD cases were however noted: in both Down syndrome and familial AD, dystrophic neurites that stained intensely for APP were frequent, but were rare in late onset AD. Such APP-positive dystrophic neurites are a long-standing finding in Down syndrome and early onset AD (Perry et al., 1988; Shoji et al., 1990; Cras et al., 1991; Joachim, et al., 1991), and it has been observed that they are relatively uncommon in late onset AD (Pera et al., 2013; Jevtic & Provia, 2019). Dissociation of TMCC2 and APP staining in familial AD may relate to the mutation of APP at residue Val717, and to an excess dose of APP in Down syndrome. Alternately this dissociation may reflect a more aggressive neuronal dysfunction and disease progression in early compared to late onset AD. All forms of AD share disrupted metabolism of APP and/or its metabolites, however mutated or excess APP is the primary driver of disease in the cases of early onset AD examined, whereas in late onset AD APOE is the principal genetic risk factor, suggesting potential differences in underlying pathomechanisms that are potentially shown by variation in the staining patterns for APP and TMCC2 despite similarities in clinical presentation.

We found an additional association of TMCC2 with AD pathology in Down syndrome dementia compared to familial AD and late onset AD. In AD associated with Down syndrome, in addition to being detected in dense cored plaques as in late onset AD, TMCC2 immunoreactivity was also found as intensities having a spicular or thread-like appearance. These prominent TMCC2- and methoxy-X04-positive, but APP-negative, features were not observed in the 10 late onset AD cases, nor in the 6 familial AD cases examined. This specific pathology of TMCC2 staining in Down syndrome may relate to over-expression of APP, its proteolytic products, or be associated with one or more of the 200-300 other genes present on chromosome 21 which will also be present in excess. In this respect, it has been noted in mouse models of trisomy 21 as well as in iPSC-derived Down syndrome neurons, that chromosome 21 genes other than APP exacerbate AD pathogenesis (Wiseman et al, 2018; Ovchinnikov et al., 2018; Sawa et al, 2021).

TMCC2 binds apoE in an apoE-isoform-specific manner, and apoE modifies an impact of TMCC2 on APP processing in cultured cells (Hopkins, et al, 2011). In age-matched non-demented and late onset AD cases stratified by APOE genotype, we found an association of APOE genotype and/or dementia with putative post-translational modification of TMCC2 (Fig. 5), suggesting an influence of APOE on the neurobiology of TMCC2 in the human brain, though an influence of dementia cannot be excluded, as we were unable to obtain age-matched non-demented APOE4 homozygotes.

Limitations of this study include its cross-sectional nature, and the number of cases examined within each of the groups investigated, which reduced statistical power for a comparison of TMCC2-related pathology with genetic influences on AD pathogenesis.

Nevertheless, across 19 cases of AD having different genetic risk profiles, we found an association of TMCC2 immunoreactivity with the pathology of AD, suggesting novel directions for investigation into the mechanisms by which TMCC2 and apoE may contribute to AD pathogenesis.

## Acknowledgements

This work was supported by an award from the MRC Confidence in Concept Kings Health Partners R & D Challenge Fund (PCRH and GT). We thank Paul Francis, Shahid Zaman and Tara Spires-Jones for advice. We thank Andrew Robinson and Federico Roncaroli at the Manchester Brain Bank, which is supported by Alzheimer’s Research UK and the Alzheimer’s Society through the Brains For Dementia Research Programme.

## Materials and Methods

### Human brain

All procedures performed in studies involving human tissue were in accordance with the ethical standards of the 1964 Helsinki declaration and its later amendments or comparable ethical studies. Ethical approval was granted by Brains for Dementia Research City and East National Research Ethics Service committee (08/H0704/128+5) and specimens provided by the Manchester Brain Bank, Salford, UK. Cases were anonymized, but information was provided regarding APOE genotype, sex, age at death, and neuropathology (supplementary Table 1).

### Western blots and Immunofluorescence

Homogenates were prepared from frozen brain tissue using an electrical homogenizer and SDS-containing buffer (4% SDS, 20% glycerol, 10% 2-mercaptoethanol, and 0.125 M Tris HCl, pH 6.8.) containing protease inhibitors (Complete, Boehringer Mannheim), followed by heating to 95°C for 5 minutes and centrifugation at 15,000 g for 10 minutes. We specifically note that efficient recovery of TMCC2 required brain homogenates to be incubated with SDS prior to centrifugation; non-ionic detergents such as Triton X-100 or NP40 did not efficiently solubilize TMCC2 from brain tissue. Rabbit anti-TMCC2 antibodies 94 and 11193 were raised against recombinant TMCC2 as previously described (Hopkins et al, 201)] and used at 1/300 for tissue staining in 10% normal goat serum, and 1/1000 for Western blots in 5% skim milk powder. SDS-PAGE gels were 10% acrylamide and loaded with 1 mg original tissue equivalent per lane. Data collection and quantification of band intensities for western blots was performed using the LI-COR Odyssey platform and statistical calculations used Graphpad Prism. Anti-Aβ antibody 4G8 was purchased from Chemicon (Temecula, CA, USA) and used at 1/300. Anti-phosphotau antibody AT8 and anti-tubulin antibody DM1A were purchased from ThermoFisher (Loughborough, UK) and used at 1/200 and 1/10,000, respectively.

For immunostaining, sections of 7µm were cut from formalin fixed paraffin embedded blocks from the same cases as used in immunoblot protocols. For chromogenic staining using 3,3’-diaminobenzidine, following dewaxing, endogenous peroxidase was blocked by 2.5% H_2_O_2_ in methanol and immunohistochemistry was performed. To enhance antigen retrieval sections were microwaved in sodium citrate pH 6.0 and kept in a citrate buffer for 10 min following heating. After blocking in normal serum (DAKO, Cambridgeshire, UK) primary antibody was applied overnight at 4 °C. Following washes, the sections were incubated with biotinylated anti-rabbit secondary antibody (DAKO), followed by avidin:biotinylated enzyme complex (Vecta-stain Elite ABC kit, Vector Laboratories, Peterborough, UK). Finally, the sections were incubated for 5–10 min with 0.5 mg/mL 3,3′-diamobenzidine chromogen (Sigma-Aldrich Company Ltd., Dorset, UK) in Tris-buffered saline (pH 7.6) containing 0.05% H_2_O_2_. For dual immunostaining, when using anti-TMCC2 (94) and AT8: following dewaxing, antigen retrieval was sodium citrate pH 6.0, 100°C for 10 minutes; for dual labeling with anti-TMCC2 (94) at 1/300, 4G8 or C1/6.1 at 1/500, this was preceded by 80% formic acid for 20 minutes. Tissue sections were blocked with 10% normal goat serum in PBS for 45 minutes and incubated with primary antibodies for 1 hour at room temperature. Secondary antibodies were Alexa-Fluor-488-conjugated goat anti-rabbit and Alexa-Fluor-546 or Alexa-Fluor-568-conjugated goat anti-mouse, from Invitrogen, and incubated for 1 hour at room temperature. Methoxy-X04 was used at 10 µM for 20 mins in PBS and autoflourescence quenched with Sudan black. Fluorescent images were collected on a Zeiss Axio Imager.Z2 LSM 800 microscope and processed using Zeiss ZEN software and ImageJ.

## Figures and Legends

**Supplementary Table S1.**
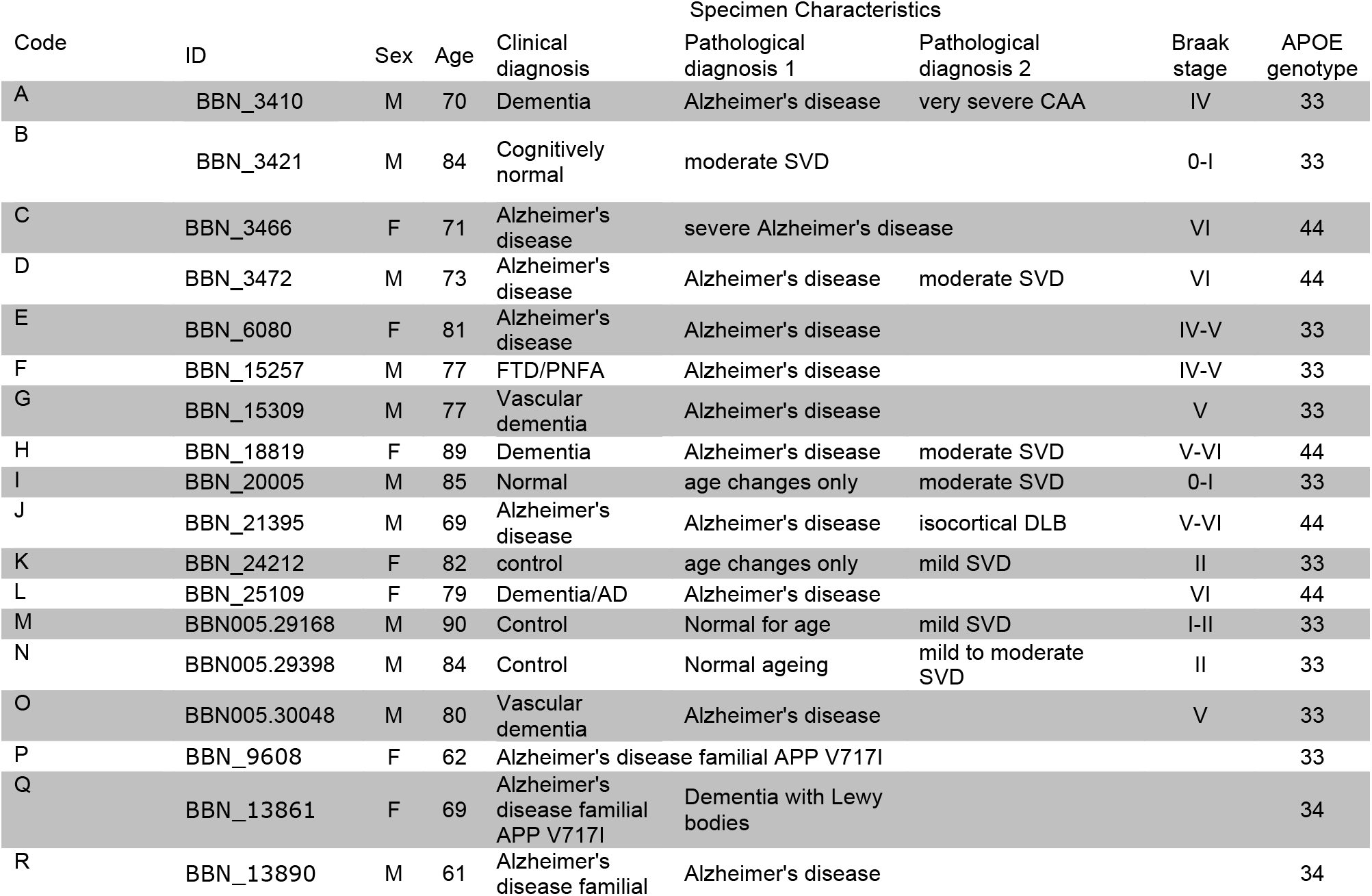
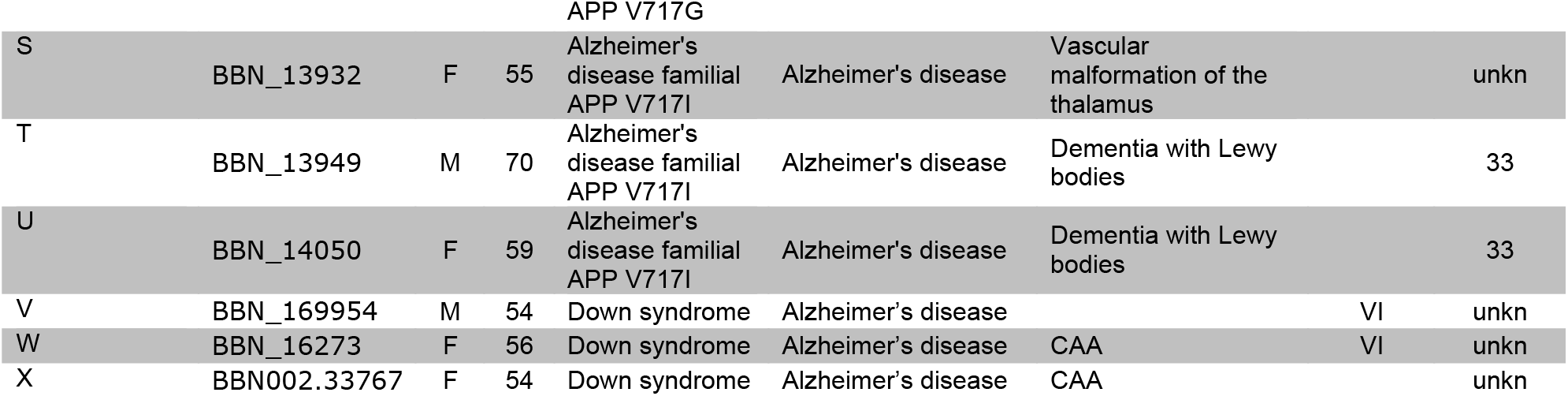
Case characteristics. AD, Alzheimer’s disease; SVD, small vessel disease; CAA, cerebral amyloid angiopathy; FTD, frontotemporal dementia; PNFA, progressive non fluent aphasia; DLB, dementia with Lewy bodies; 33, homozygous for APOE3; 44, homozygous for APOE4; unkn, unknown.

## Supplementary Figures

**Figure S1.**
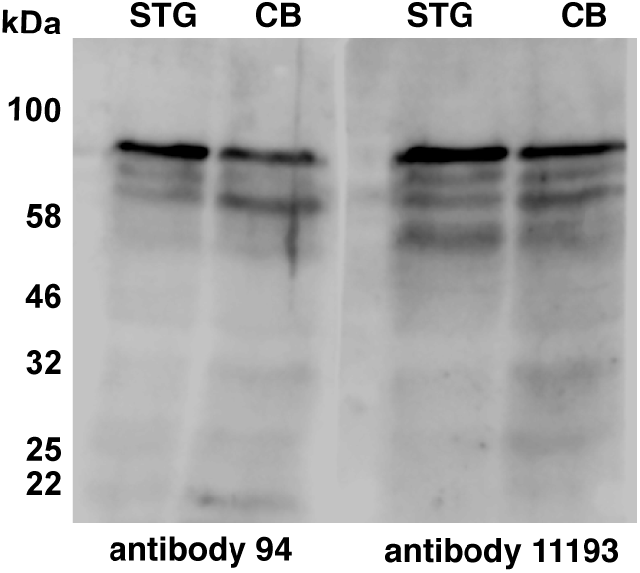
Western blot of human temporal cortex or cerebellum pools for TMCC2 using rabbit anti-TMCC2 antibody 11193 confirming specificity of the TMCC2 and TMCC2* bands detected by antibody 94 shown in Fig. 1A and Fig 5.

**Figure S2.**
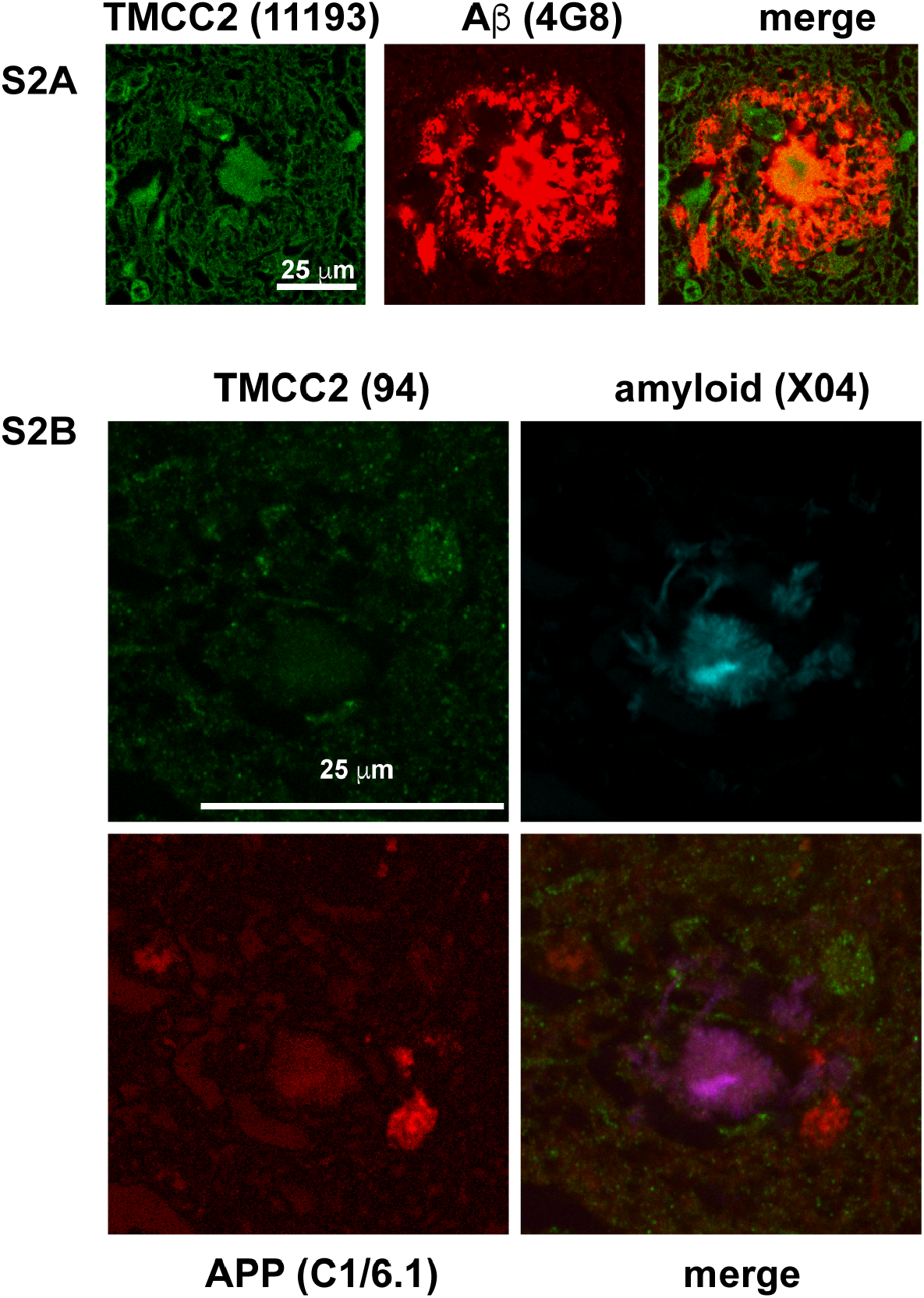
**S2A**, detection of TMCC2 using rabbit anti-TMCC2 antibody 11193 in dense-cored amyloid plaques identified using anti-Aβ antibody 4G8. **S2B**, example of uncommon separate staining for TMCC2 and APP in dystrophic neurites adjacent to dense cored amyloid plaques in late onset AD.

**Figure S5.**
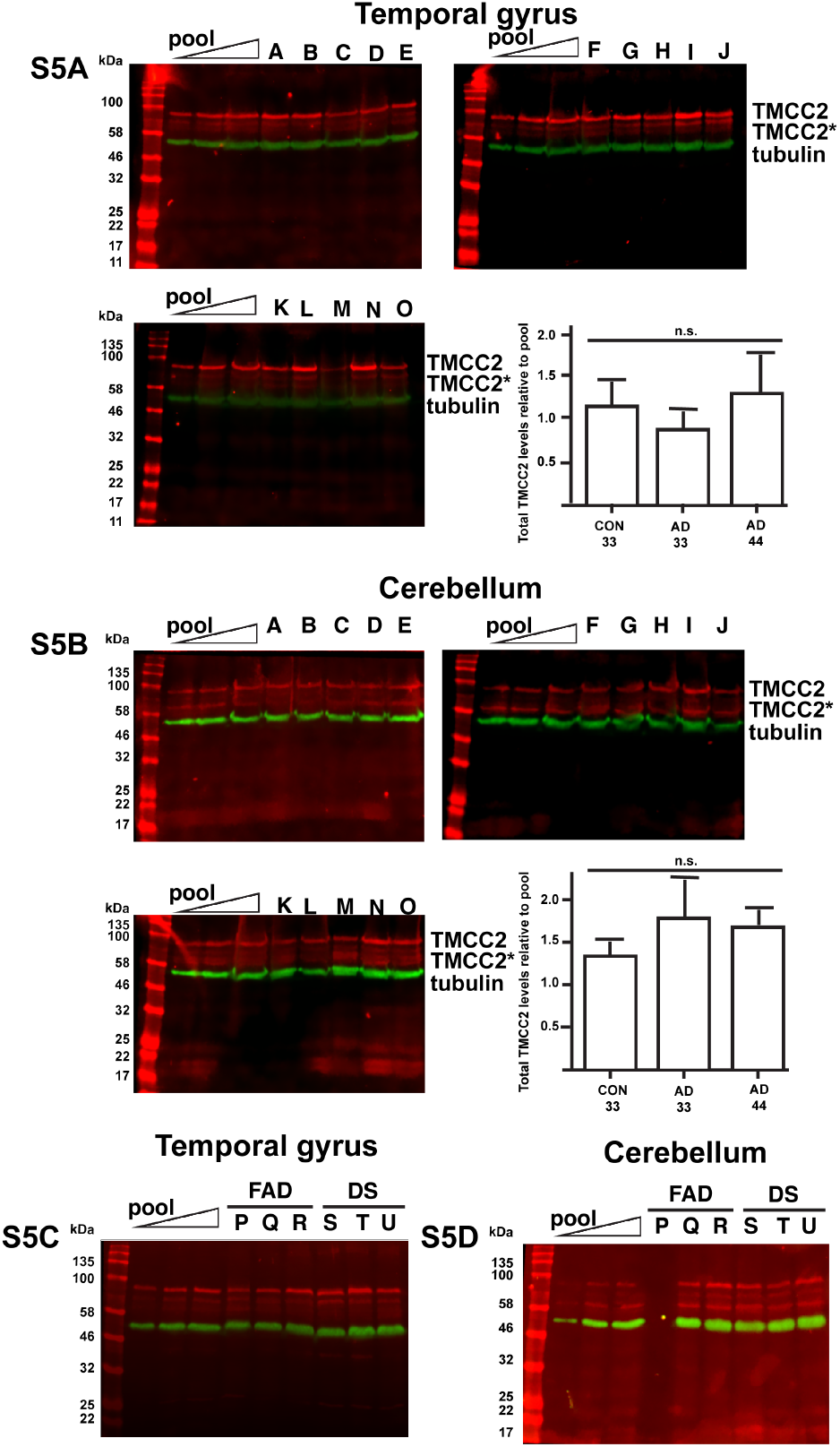
Alteration of TMCC2 according to brain region. **S5A**, western blot and quantification of total TMCC2 levels from human temporal cortex using antibody 94. **S5B**, western blot and quantification of total TMCC2 levels from human cerebellum using antibody 94. Pools were loaded at a ratio of 0.5 to 1 to 2, with 1 representing approximately 1 mg of original tissue. Bar charts show the relative levels of total TMCC2 for control APOE3 homozygotes, and AD cases homozygous for either APOE3 or APOE4 normalized between blots by reference to the band intensities of the pools; no significant association of APOE genotype or AD with total TMCC2 levels was found (p=0.63 for temporal gyrus and 0.24 for cerebellum, 1 way ANOVA). **S5C** and **S5D**, western blot of temporal gyrus and cerebellum, respectively, from early onset AD cases analysed as above. Frozen tissue from the cerebellum of case P was not available. Letters above lanes refer to cases in Table S1.

